# Modeling visual performance differences with polar angle: A computational observer approach

**DOI:** 10.1101/434514

**Authors:** Eline R. Kupers, Marisa Carrasco, Jonathan Winawer

## Abstract

Visual performance depends on polar angle, even when eccentricity is held constant; on many psychophysical tasks observers perform best when stimuli are presented on the horizontal meridian, worst on the upper vertical, and intermediate on the lower vertical meridian. This variation in performance ‘around’ the visual field can be as pronounced as that of doubling the stimulus eccentricity. The causes of these asymmetries in performance are largely unknown. Some factors in the eye, e.g. cone density, are positively correlated with the reported variations in visual performance with polar angle. However, the question remains whether such correlations can quantitatively explain the perceptual differences observed ‘around’ the visual field. To investigate the extent to which the earliest stages of vision –optical quality and cone density- contribute to performance differences with polar angle, we created a computational observer model. The model uses the open-source software package ISETBIO to simulate an orientation discrimination task for which visual performance differs with polar angle. The model starts from the photons emitted by a display, which pass through simulated human optics with fixational eye movements, followed by cone isomerizations in the retina. Finally, we classify stimulus orientation using a support vector machine to learn a linear classifier on the photon absorptions. To account for the 30% increase in contrast thresholds for upper vertical compared to horizontal meridian, as observed psychophysically on the same task, our computational observer model would require either an increase of ~7 diopters of defocus or a reduction of 500% in cone density. These values far exceed the actual variations as a function of polar angle observed in human eyes. Therefore, we conclude that these factors in the eye only account for a small fraction of differences in visual performance with polar angle. Substantial additional asymmetries must arise in later retinal and/or cortical processing.

**Author Summary:** A fundamental goal in computational neuroscience is to link known facts from biology with behavior. Here, we considered visual behavior, specifically the fact that people are better at visual tasks performed to the left or right of the center of gaze, compared to above or below at the same distance from gaze. We sought to understand what aspects of biology govern this fundamental pattern in visual behavior. To do so, we implemented a computational observer model that incorporates known facts about the front end of the human visual system, including optics, eye movements, and the photoreceptor array in the retina. We found that even though some of these properties are *correlated* with performance, they fall far short of *quantitatively explaining it*. We conclude that later stages of processing in the nervous system greatly amplify small differences in the way the eye samples the visual world, resulting in strikingly different performance around the visual field.

## 1. Introduction

### 1.1 Psychophysical performance differs with visual field position

Psychophysical performance is not uniform across the visual field. The largest source of this non-uniformity is eccentricity: acuity is much higher in the central visual field (fovea), limiting many recognition tasks such as reading and face recognition to only a relatively small portion of the retina. As a result, central visual field loss, such as macular degeneration, can be debilitating. Even a modest difference in eccentricity can have substantial effects on performance. For example, contrast thresholds on an orientation discrimination task approximately triple at 8° compared to 4° eccentricity (1, 2). Similar effects are found for a wide range of tasks (for a review on peripheral vision, see (3)).

Interestingly, visual performance differs not only as a function of distance from the fovea (eccentricity), but also around the visual field (polar angle). The polar angle effects can be quite systematic. At a fixed eccentricity, contrast sensitivity and spatial resolution are better along the horizontal than the vertical meridian, and better along the lower than the upper vertical meridian. The two effects have been described by Carrasco and colleagues and called the “horizontal-vertical anisotropy” and the “vertical meridian asymmetry” (2, 4–8). Such effects, often called *performance fields*, are found in numerous tasks, including contrast sensitivity and spatial resolution (5, 6, 9–23) (**Fig 1**), visual search (24–31), crowding (32–34), motion perception (35), visual short-term memory (36), contrast appearance (7) and spatial frequency appearance (37). These effects can be large. For example, contrast thresholds at 8° eccentricity can be 5 times lower on the horizontal meridian compared to the vertical (5), a larger effect than doubling the eccentricity, from 4° to 8° (2).

**Fig 1.**
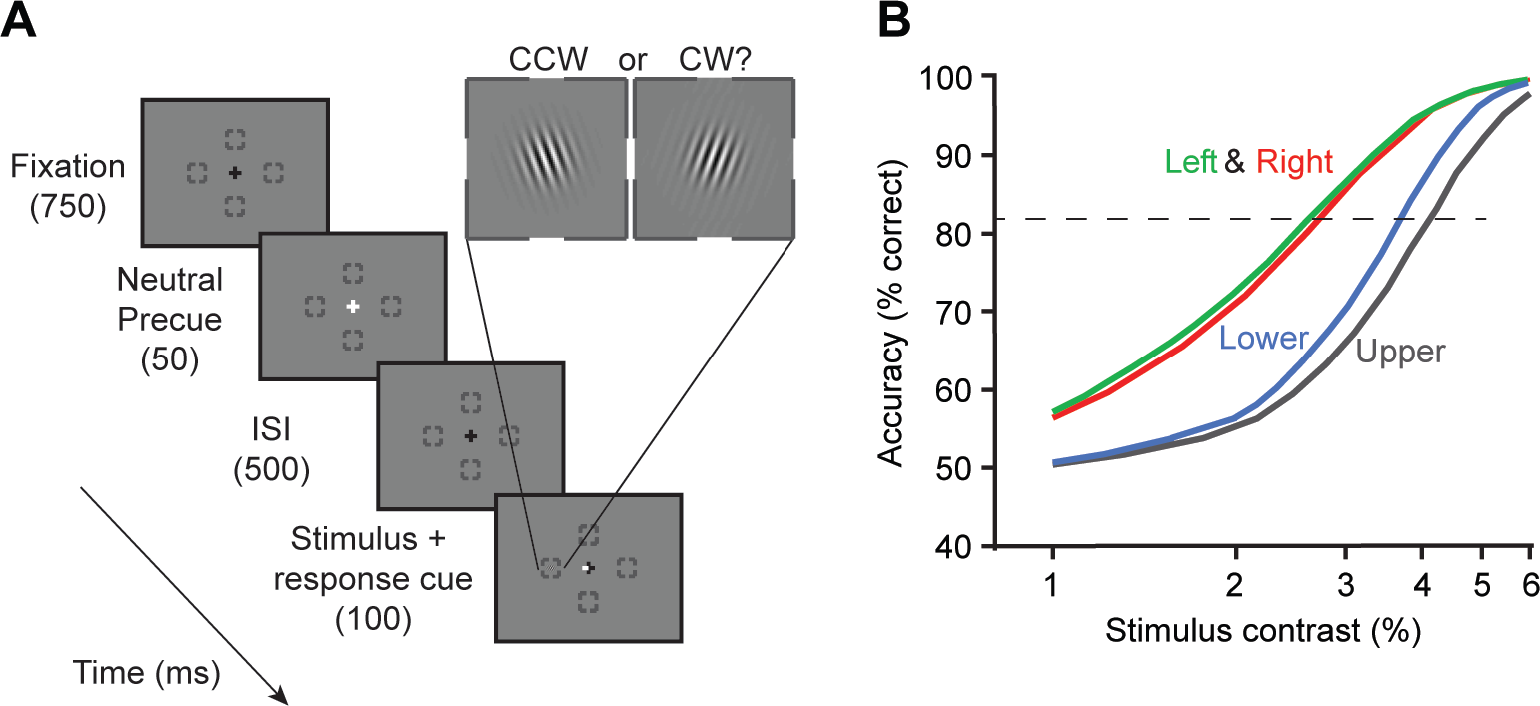
Example of psychophysical task and performance differences ‘around’ the visual field. **(A)** Experimental design of a two-alternative forced choice (2-AFC) orientation discrimination task. While observers maintained fixation, a brief cue appeared, and after a 500 ms ISI and after a Gabor stimulus (4 cycles/°) was presented at one of four possible iso-eccentric locations (4.5° eccentricity). A response cue indicated to the observer the target location they were asked to make an orientation judgment on (clockwise/CW or counter-clockwise /CCW relative to vertical). Contrast varied across trials. **(B)** Contrast-dependent psychometric functions of an example observer show best performance for the two horizontal locations (Left and Right), and poorest performance for upper vertical. The increase in contrast thresholds (contrast at which performance reached 82% correct, dashed line) ranged from about 2.5% (Left and Right) to 4% (Upper). Data from figure 6 in Cameron, Tai & Carrasco (5).

The causes of performance differences with polar angle are not known. Some eye factors may contribute to them as they do to differences across eccentricity. For instance, the drop-off in density of cones and retinal ganglion cells with eccentricity contributes to decreased acuity (38, 39). In this paper, we take a modeling approach to quantify the extent to which optics and photoreceptor sampling contribute to performance differences with polar angle.

### 1.2 Cone density differs with visual field position

In the human eye, cone density varies with eccentricity and polar angle. Foveal cones have relatively small diameters and are tightly packed, becoming sparser in the periphery due to both increased size and larger gaps between them (40, 41).

Cone density also differs as a function of polar angle at a fixed eccentricity. From ~2° to 7° eccentricity, density is about 30% greater on the horizontal than the vertical meridian (40, 41) (**Fig 2**). This 30% difference is about the same as the cone density decrease from 3° to 4° eccentricity along a single meridian. As a result, iso-density contours are elongated by about 30% in the vertical axis compared to horizontal. Because cone density is higher on the meridian where performance is better (horizontal compared to vertical), one might be tempted to conclude that cone density explains the performance difference. We return to this question in subsequent sections.

**Fig 2.**
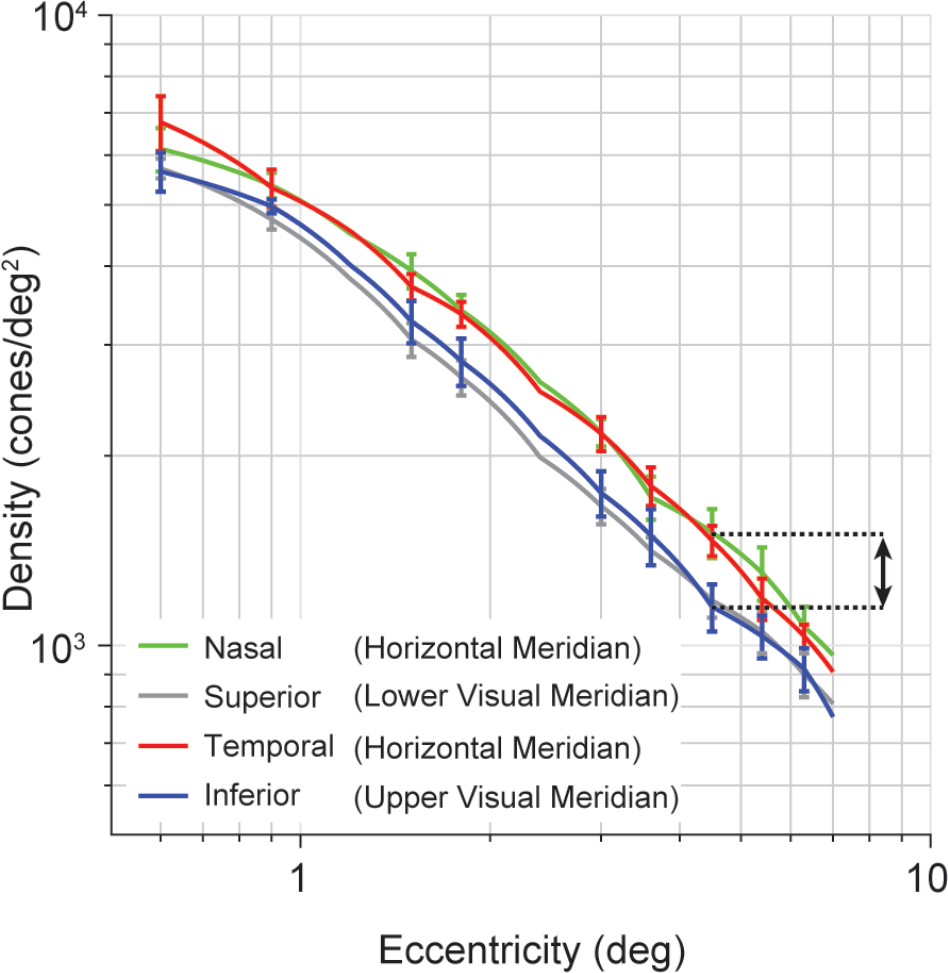
Variations in human cone density as a function of eccentricity and polar angle. Cone density falls off sharply with eccentricity, and also varies between horizontal and vertical retinal axes. Data are pooled across both eyes. Error bars represent standard error across 10 observers. Arrow indicates difference in cone density between horizontal and upper visual meridian at 4.5° eccentricity (matching **Fig 1**). Figure recomputed with data from observers between 22-35 years old (‘Group 1’) reported by Song et al. (41).

### 1.3 Optical quality differs with visual field position

Before light hits the retina, it has already been transformed by refraction from passing through different media (cornea, vitreous and aqueous humors), by diffraction from the pupil, as well as by optical aberrations (chromatic and achromatic) of the lens and intraocular light scattering (for an overview see (42)). These transformations reduce the optical quality of the image projected onto the retina.

Optical quality is not uniform across the retina (43, 44). A clear, systematic effect is that both defocus and higher-order aberrations become worse with eccentricity. We assume that myopes and hyperopes wear corrective lenses to achieve good focus at the fovea. Defocus is the largest contribution to image quality (42). The effects of defocus are largest in the far periphery, but still evident in when comparing fovea to parafovea (e.g., 0 vs 5°, **Fig 3**).

**Fig 3.**
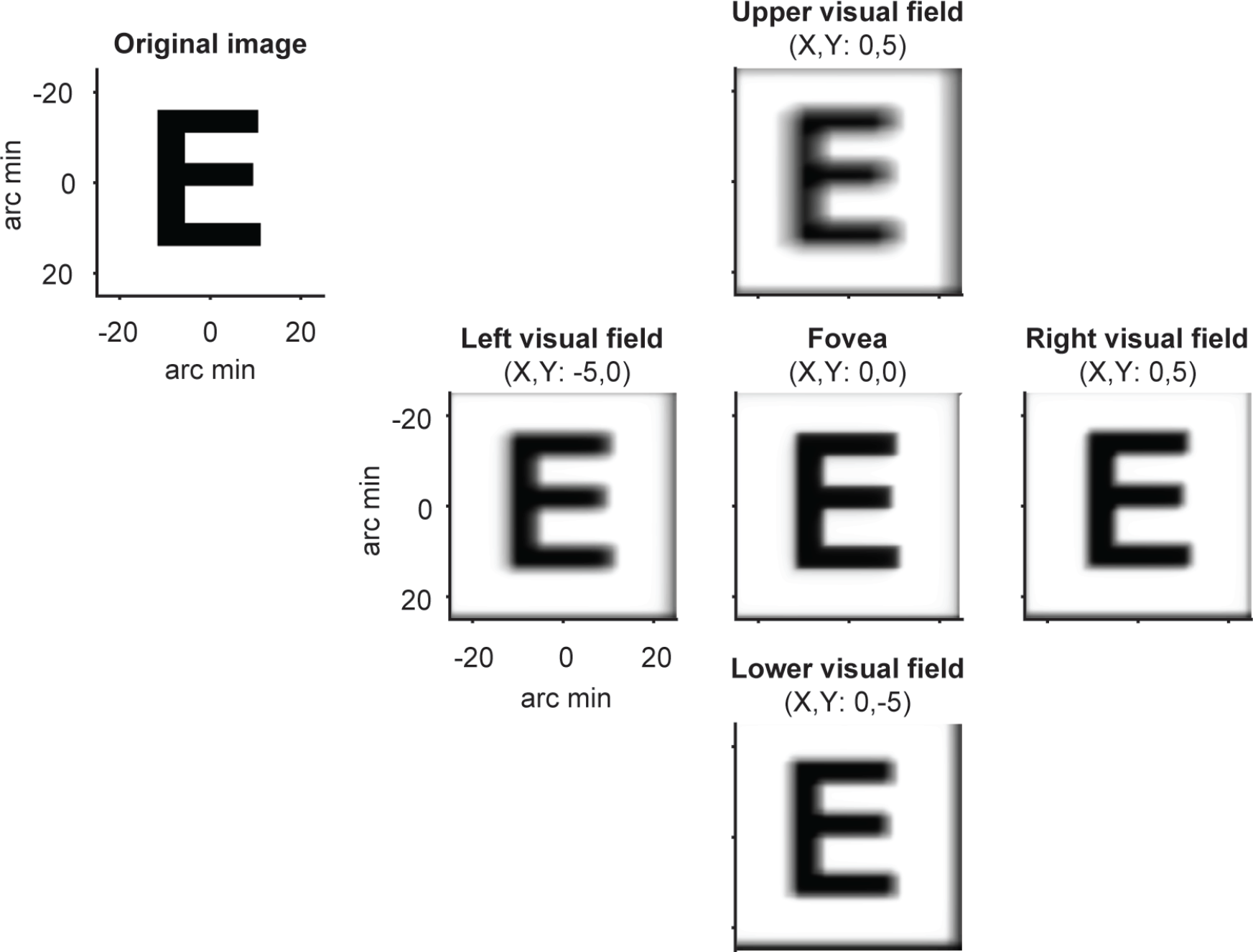
Variations in optical quality as a function of visual field location for example observer. The letter E was convolved with 5 location-specific point spread functions (PSFs) from an example observer, using wavefront measurements of image quality and correcting for the central refractive error. The wavefront measures are based on a prior study (43), and provided courtesy of Pablo Artal.

Most measurements of optical quality in human are either at the fovea or along the horizontal meridian. These measurements show that in addition to the decline in optical quality with eccentricity, there are also hemifield effects: For example, in the periphery, the temporal retina tends to have poorer optics than the nasal retina (44). There are some (43), but many fewer measurements along the vertical meridian compared to the horizontal meridian. To our knowledge, it is not yet firmly established whether there are systematic differences in optical quality between the vertical and horizontal meridians. However, the fact that optical quality varies with eccentricity as well as between nasal and temporal retina suggests that one should at least consider optics as a possible explanatory factor for performance differences around the visual field.

### 1.4 Quantifying the contribution of components in the eye to behavioral performance using a computational observer model

The meridian differences in cone density are correlated with meridian differences in psychophysical performance, with higher cone density and better performance on the horizontal axis (adjusted R^2^ = 0.88, **Fig 4**) (40, 41). However, a correlation does not necessarily imply an explanation. Without an explicit linking hypothesis or model that can predict how a difference in cone density should affect visual performance on a given task, we cannot know whether this correlation is meaningful for explaining this behavior. Should a decrease in cone density increase contrast thresholds? And if so, should a decrease of about 25% in cone density (difference between horizontal and vertical at 4.5° eccentricity) lead to an increase in contrast threshold of about 25%, as observed psychophysically?

**Fig 4.**
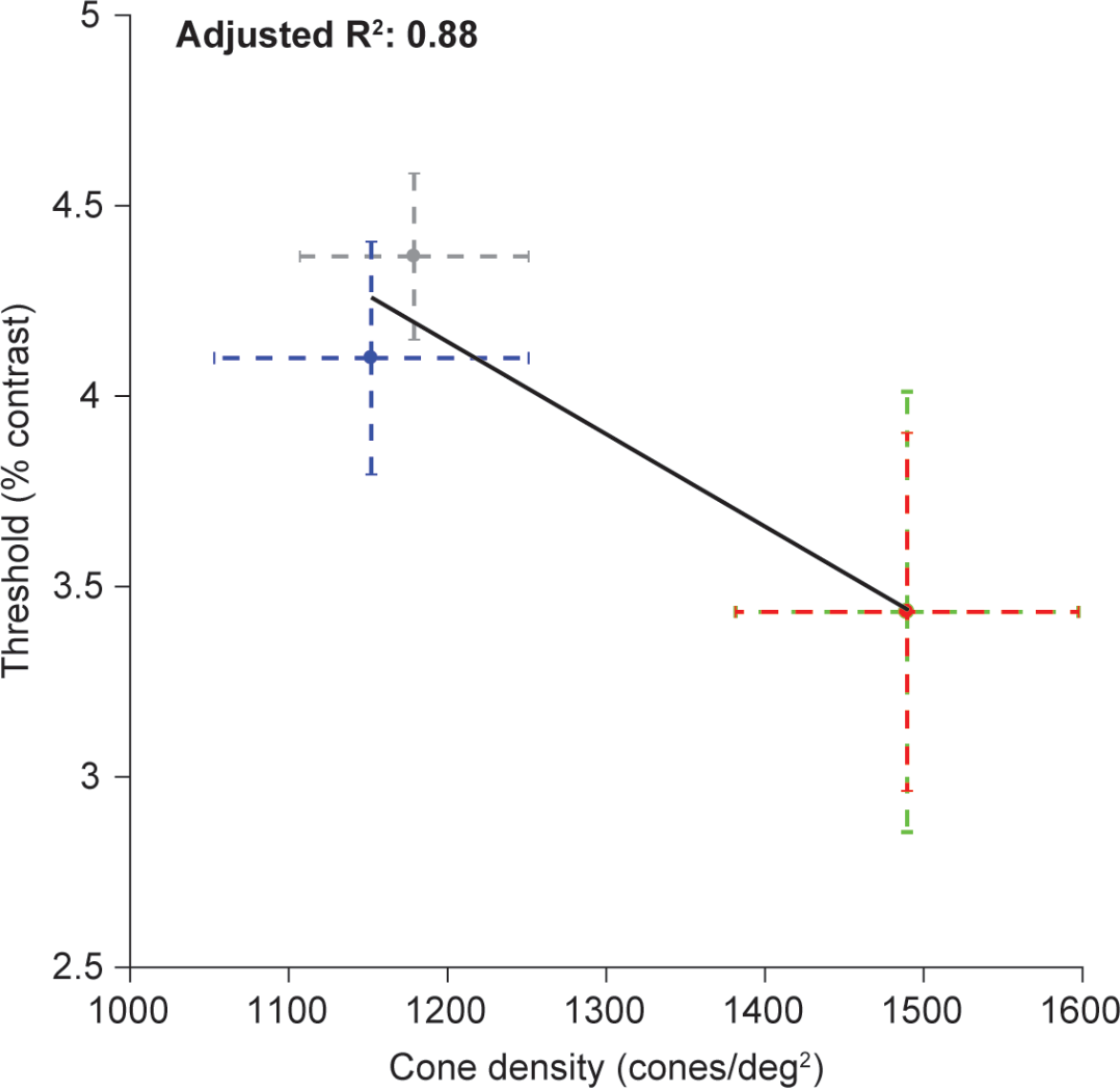
Correlation between performance and cone density across polar angles. Contrast thresholds (y-axis) were averaged across three observers reported in Cameron, Tai, and Carrasco (5). Thresholds were obtained for stimuli at 4.5° eccentricity, above (gray), below (blue), left (red), and right (green) of fixation. Cone density along the four meridians are the average values reported by Song et al. (41) at 4.5° eccentricity. (For right and left, cone density was averaged for the nasal and temporal retina, since observers performed the psychophysics task binocularly). Error bars indicate one standard error across 10 observers (cone density) and 3 observers (contrast thresholds).

Answering these questions requires a computational model. A computational model of the eye can quantify the extent of each component’s contribution on visual performance and potentially reveal which components limit performance on a given task.

In this study, we quantify the contribution of cone density and optical quality on visual performance according to a computational observer model. We then compare the modeled contributions to the observed quantities, and ask whether the observed differences in cone density and optical quality as a function of polar angle can explain the observed differences in performance.

To implement the computational observer model of the human eye, we used the Image Systems Engineering Toolbox for Biology (ISETBIO (45–47)), a publicly available toolbox, to simulate encoding stages in the front-end of the human visual (available at http://isetbio.org/). We used this model to simulate a 2-AFC orientation discrimination task using Gabor stimuli matched in parameters as reported by Cameron, Tai and Carrasco (5). Our computational observer model consists of multiple stages representing the front-end of the visual system: the spectral radiance of the experimental visual stimuli, optical quality of the lens and cornea, fixational eye movements, the cone mosaic with photoreceptors and their isomerization rate for a given stimulus presentation.

The goal of a computational observer model is to calculate the effect of a stimulus manipulation at particular encoding stages of the visual pathway, where each stage is represented as biologically plausible as possible and including sources of noise. In our model, we include photon noise, noise in the visual system and uncertainty about the decision when discriminating between two stimulus classes. Given that our computational observer model does not have access to all the information of the stimulus when executing the task and must learn the stimulus class from noisy data, it will <u>not</u> represent optimal performance for the given task. Our model is therefore different from ideal observer models: i.e. models that have access to all information when executing the task and quantify optimal performance at a given encoding stage because it is only limited by biophysical constraints.

With a computational observer model, one can show where along the visual pathway information loss happens and how this loss of information is inherited or potentially compensated for in later encoding stages of the visual pathway. Here, we investigate to what extent visual performance (contrast threshold) depends on variations in cone density and optical quality. By systematically varying cone density and optical quality (defocus) independently, we can compare the computational observer model performance to reported differences in the literature on the human eye to quantify the individual contribution of each of these two factors to differences in visual performance across the visual field.

## 2. Results

### 2.1 Overview of computational observer model

To investigate to what extent performance differences in the visual field depend on variations in cone density and optical quality, we developed a computational observer model of the first stages of the human visual pathway. The computational observer was presented with oriented Gabor stimuli, tilted either clockwise or counter-clockwise from vertical, to simulate a 2-AFC orientation discrimination task. To compare the performance of the computational observer to human observers, we matched the stimulus parameters to a psychophysics study (5).

#### 2.1.1 Scene radiance

The model starts from the photons emitted by a visual display, defined as the scene radiance. Three frames of two example time-varying achromatic Gabor stimuli are shown in the first panel of **Fig 5**. The left column shows the result of each stage for a Gabor with high contrast (100% black outline) and middle column for a low contrast Gabor (10%, orange outline). Both Gabor stimuli are oriented clock-wise from vertical, contain a spatial frequency of 4 cycles/° and are presented at 4.5° eccentricities. Photons were emitted from pixels whose spectral power distributions matched those of an Apple LCD display. The high and low contrast Gabor stimuli contained the same mean radiance, but the high contrast Gabor has an amplitude that is 10x larger than the low contrast Gabor (1D representation at dotted line, right column of panel 1).

**Fig 5.**
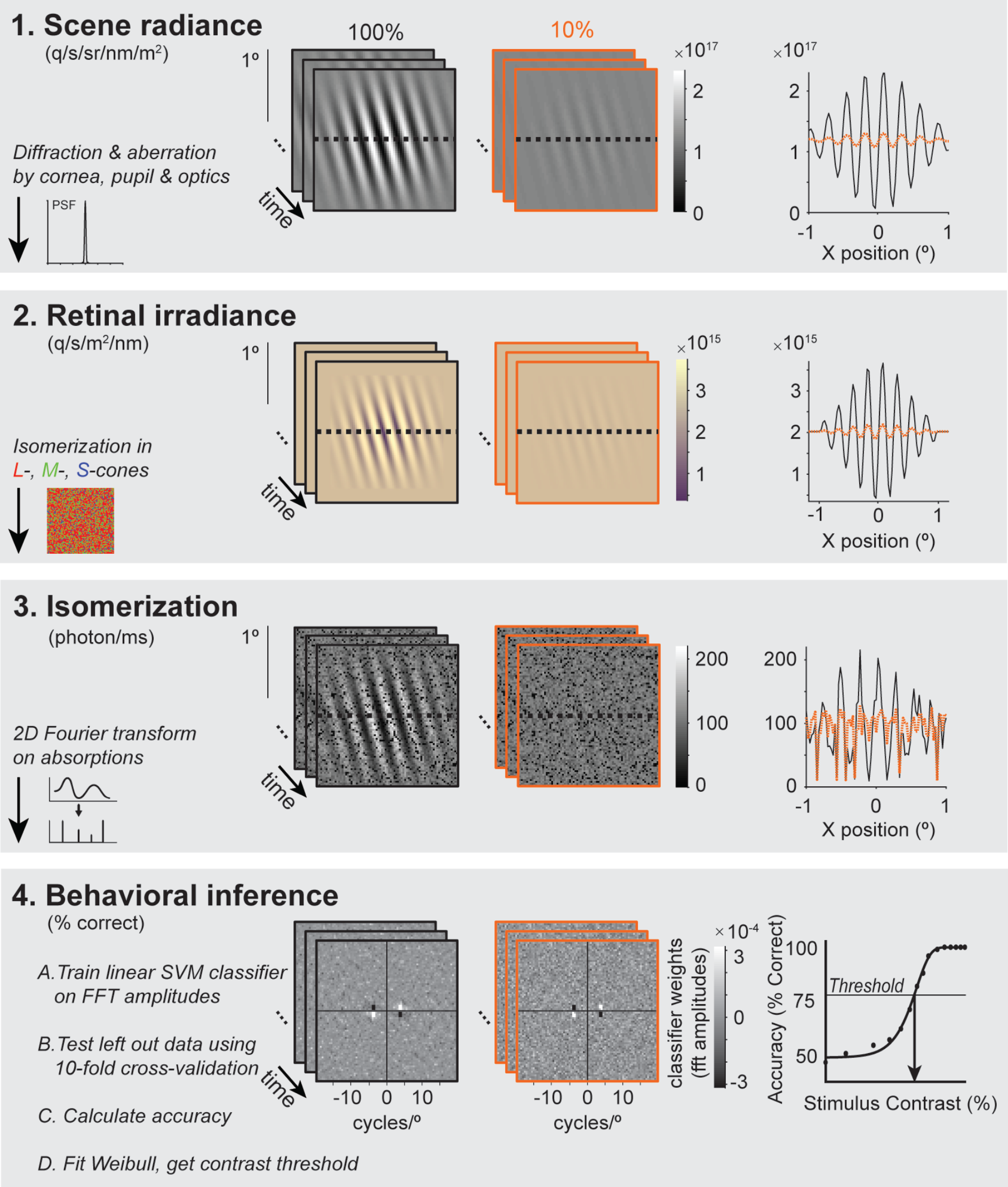
Overview of computations in observer model. Computations are shown for a 100% (black outline) or 10% (orange outline) contrast Gabor stimulus. **(1) Spectral radiance.** The computation starts with the spectral radiance of a time-varying scene. The 2D images shows the radiance summed across wavelengths (400-700 nm) during 1-ms windows. 1D slices through the images are plotted on the right. **(2) Retinal irradiance.** The retinal irradiance is the image that has passed through the eye’s optics, before processed by the retina. The optics are modeled as a typical human wavefront with a 3-mm pupil, indicated by the schematic point spread function (PSF) on the lower left. For illustration purposes, wavelengths of the 2D representation were converted into RGB values. The yellowing is a result of the spectral filtering by the eye’s optics. **(3) Isomerization.** The time-varying retinal irradiance is transformed into photon absorptions using a rectangular grid of L-, M- and S-cones. The dark pixels correspond to S-cones (see text for more details). **(4) Behavioral inference.** The last stage of the computational observer is a simplified decision stage, performing a 2-AFC orientation discrimination task. First, the absorption image at each time point was transformed into the amplitude domain by the Fourier transform. The amplitude images were used to train a linear support vector machine (SVM) classifier using 10-fold cross-validation. The images show an example of the trained classifier weights. The weights are high at locations corresponding to the stimulus frequency (4 cycles/°) and orientation (±15°). Once trained, the classifier predicted the left-out data. The cross-validated accuracy was fit by a Weibull function (right panel) to determine the contrast threshold.

#### 2.1.2 Retinal irradiance

The second stage of the model simulates the retinal irradiance: the result of the time-varying radiance passing through the simulated human optics (including refraction and aberrations caused by the pupil, cornea, and lens). The retinal irradiance is the light image just before the photons are captured by the photopigment in the retina. The second panel of **Fig 5** shows the retinal irradiance summed across all wavelengths. The effect of the optics is to blur the Gabor stimuli and to reduce the fraction of short wavelength light. The mean irradiance is the same for the two stimuli.

#### 2.1.3 Isomerization

The third stage implements a cone mosaic and computes photon absorptions for each cone at each time sample (panel 3 of **Fig 5**). The cone mosaic is a rectangular patch with a field of view of 2×2° at 4.5° eccentricity. Each cone type absorbs a percentage of the emitted photons, depending on the wavelength of the light and the efficiency of the cone type.

The model implements two sources of noise. The first source comes from small fixational eye movements. These eye movements cause shifts of the stimulus on the cone mosaic during the trial. The second noise source is from photons, which are inherently noisy and follow a Poisson distribution.

During the onset of a high contrast Gabor, the L-, M- and S-cones increase their absorptions on average by ~50, ~30 and ~4 photons/ms respectively. After stimulus offset, the absorptions return to baseline at ~110, ~75 and ~12 photons/ms. The absorption rates for a mean luminance screen (~110 photons/ms for the L-cones) was validated by an independent computation of isomerization given the luminance given by Wyszecki and Stiles (48), implemented in the ideal observer model by Geisler (equation 2, p.776, (49)), where the average L-cone absorption under 100 cd/m^2^ with a 3-mm pupil is predicted as ~108 photons/ms. The S-cones absorb fewer photons than the L- and M-cone. This is because inert pigments in the lens and macula absorb more light at short wavelengths, and because the photopigment density is lower in the S-cones. As a result, the locations of S-cones in absorption array (**Fig 5**, panel 3) are dark.

#### 2.1.4 Behavioral inference

The last stage computes a 2-AFC inference of the stimulus orientation from the cone absorptions. An ideal observer would have full knowledge of the response statistics (mean and distributions of responses of each cone at each time point from each of the possible stimuli), and uses this full knowledge to make the optimal decision given a particular measurement. It is unlikely that human decision-making has access to this full knowledge. Here we implement a computational observer which learns patterns from the data, using a linear support vector machine (SVM) classifier. The classifier uses the weights learned from training data to classify the stimulus orientation of the left-out data.

Because the within-class stimuli differ in phase and because the eyes move during the trial, the outputs of individual cones are not informative about the decision. Hence a linear classifier trained directly on the cone outputs would fail. We therefore transform the cone outputs prior to training the classifier by computing the 2D Fourier transform on the cone array at each time point. We retain the amplitudes and discard the phase information. Because the Fourier transform separates the phase and amplitude for each spatial frequency and orientation, with sufficient signal to noise it is possible to infer the stimulus orientation (irrespective of phase) from the amplitude spectrum. Transforming the outputs of the cone array in this way can be thought of as giving the observer model partial information about the task: namely, that orientation and spatial frequency (but not phase) might be relevant.

As a proof of concept, the classifier shows two expected patterns. First, the largest weights of the classifier are centered on the peak spatial frequency (4 cycles/°) and orientations (±15°) of the stimuli (**Fig 5**, panel 4). Second, the classifier accuracy increases with stimulus contrast (**Fig 5**, panel 4, right). To summarize the data for a given simulated experiment, we computed the contrast threshold by fitting a Weibull function to the cross-validated accuracy as a function of stimulus contrast. The contrast threshold for the computational observer with typical human optics and a cone mosaic matched to ~4.5° eccentricity was 2.7%. This is slightly lower (thus better performance) than thresholds reported in the psychophysics experiment with the same stimulus parameters (5), which ranged from 3.6-9% contrast for human observers.

### 2.2 The effect of small fixational eye movements in the computational observer model

Our computational observer model includes small fixational eye movements (**Fig 6A**). We implement drift based on a statistical model by Mergenthaler and Engbert (50) and microsaccades based on statistics reported by Martinez-Conde et al. (51, 52). The displacement of the stimulus due to eye movements in our simulations is relatively small: within one trial (a single colored line), the retinal displacement tends to be about 2-4 cones or less (**Fig 6A**). This is small compared to the spatial scale of our stimulus, for which a full cycle corresponds to ~6 cones at 4.5° eccentricity. Given that the trials last only 54 ms, the probability of a microsaccade is low. Hence when both microsaccades and drift are present, eye movements are dominated by drift.

**Fig 6.**
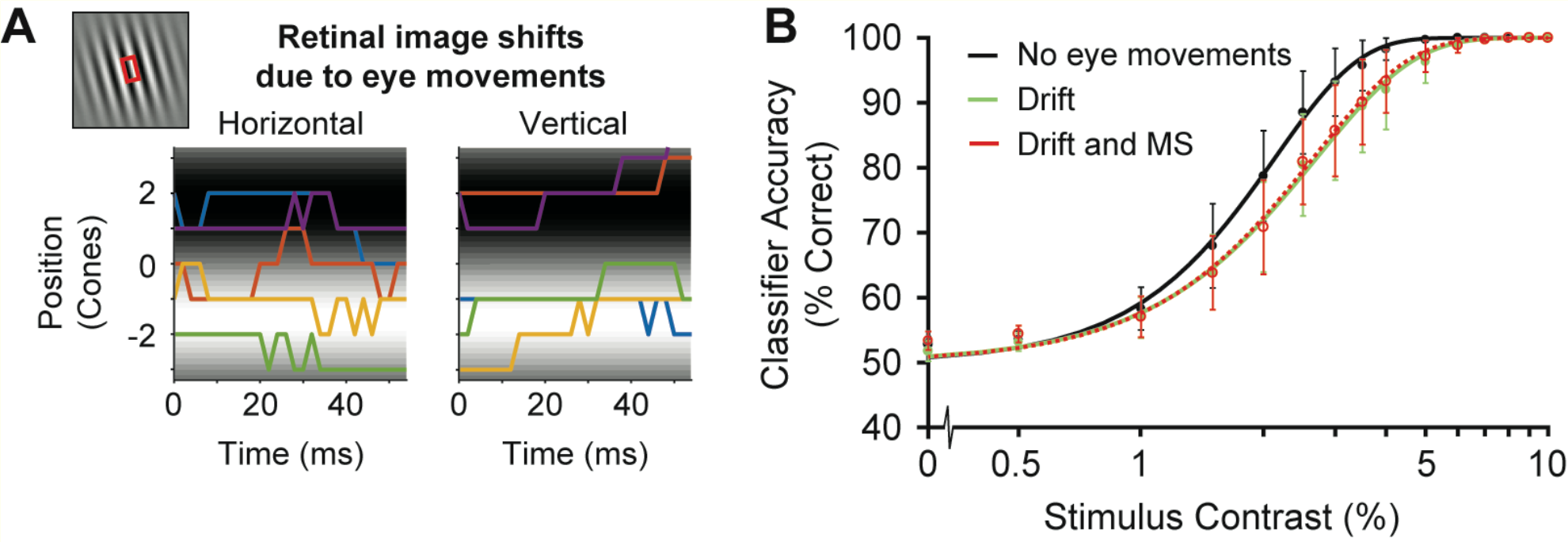
The effect of fixational eye movements on model performance. **(A)** Horizontal and vertical and 2D displacements in units of cones of the retinal image over time. Colors indicate eye movement paths of example 5 trials. A 1D slice of a single cycle of the Gabor stimulus is shown as the background (red box). **(B)** Computational observer performance with no eye movements (black), only drift (green) or both drift and microsaccades (red). Accuracy is averaged across performance of 2,000 trials per stimulus contrast (30,000 trials in total). Error bars represent standard error of the mean across 5 simulated experiments with 400 trials for each stimulus contrast.

The fixational eye movements have a small but systematic effect on the computational observer model, making performance slightly worse (**Fig 6B**). It might be surprising that eye movements have any effect on the model performance: An image translation is equivalent to a phase shift in the Fourier domain and the model discards phase information. However, because the retinal mosaic contains multiple cone types with different sensitivities, a shift in the stimulus causes a change in both the amplitude and phase spectra of the absorption images, affecting the information available to the classifier.

### 2.3 The effect of optical quality on orientation discrimination

Large levels of defocus worsen visual acuity (53), where defocus levels larger than 0.75 diopters (corresponding to 20/40 vision on the Snellen acuity chart for near sightedness) are usually compensated for with visual aids. Here, we tested the effect of defocus on the 2-AFC orientation discrimination task reported by Cameron, Tai and Carrasco (5) to investigate whether variations in defocus could explain the decrease in performance with polar angle. If the task is very sensitive to the level of defocus, then small differences in optical quality as a function of polar angle might explain the observed differences in performance.

Defocus affects the modulation transfer function of a typical human wavefront by attenuating the high frequencies (**Fig 7A**). The Gabor patches in our experiment had a peak spatial frequency of 4 cycles/° (dashed line). For this spatial frequency, the simulated levels of defocus in the observer model cause a modest reduction in contrast.

**Fig 7.**
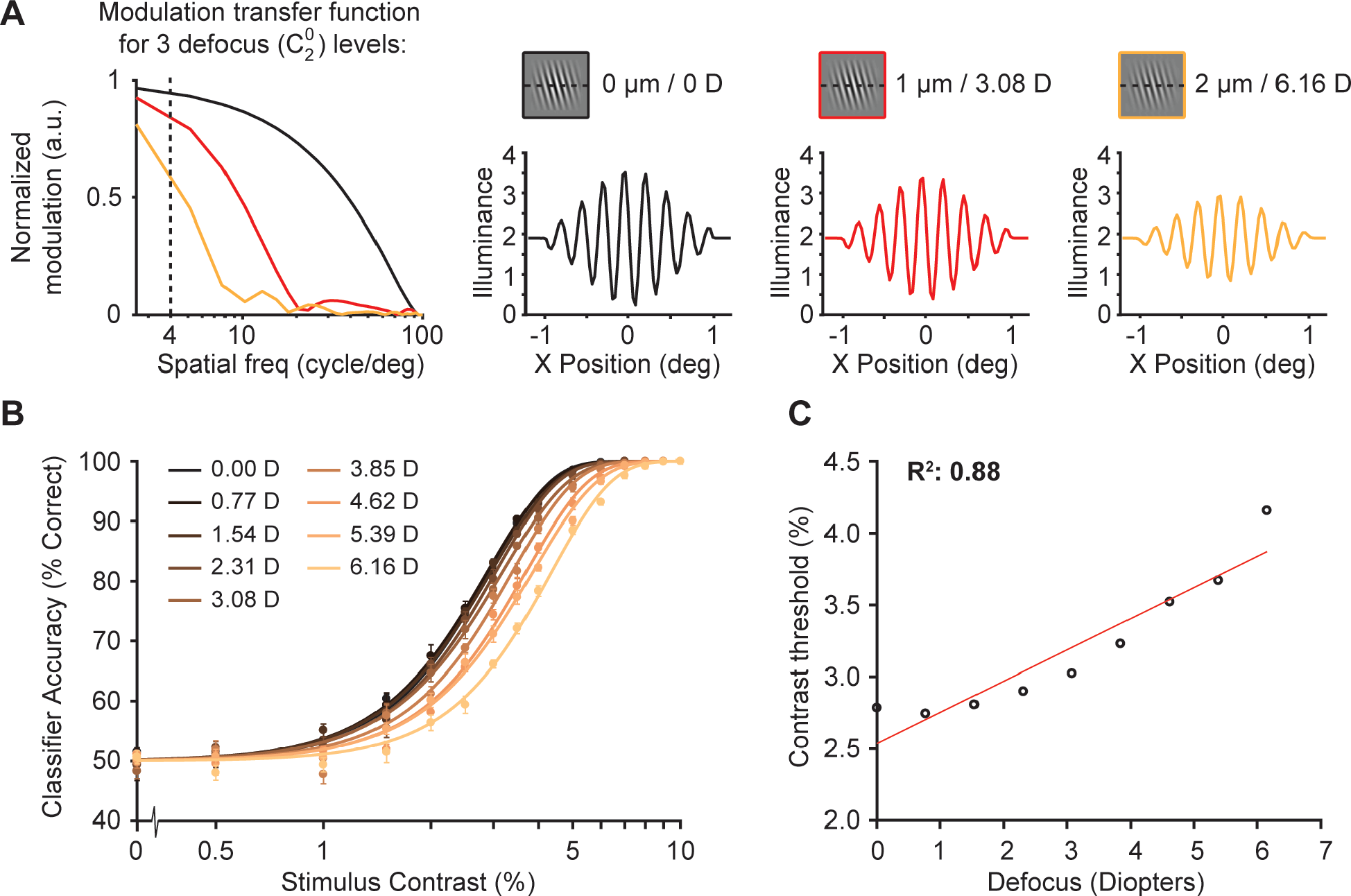
The effect of optical quality on model performance. **(A)** The modulation transfer function (MTF) of our computational observer model is shown using different levels of defocus (an addition to diffraction and higher order aberrations). The MTFs are based on a typical human wavefronts using the statistical model provided by Thibos (54) based on data from Thibos et al. (55). The three example MTFs in black, red and yellow lines indicate a defocus Zernike coefficient of 0, 1 or 2 μm respectively (equivalent to 0, 3.08 or 6.16 diopters for a 3-mm pupil). Dotted line represents 4 cycles/°. A 1D slice through the stimulus shows the effect of the three levels of defocus. **(B)** Classifier accuracy for absorption rates as a function of stimulus contrast for 9 different defocus levels for a cone mosaic simulated at 4.5° eccentricity. Accuracy is averaged across performance of 2,000 trials per stimulus contrast (30,000 trials in total). Error bars represent standard error of the mean across 5 simulated experiments with 400 trials for each stimulus contrast. **(C)** Contrast thresholds from panel B as a function of defocus in diopters, fitted with a linear function.

As expected, large increases in defocus cause the computational observer model to perform worse, evidenced by a rightward shift of the psychometric curve (**Fig 7B**). When comparing contrast thresholds as a function of defocus level, the computational observer model shows an approximately linear relation between contrast threshold and defocus (R^2^ = 0.88, **Fig 7C**). However, the effect of defocus on model performance is small. To explain an increase of 1.5% in contrast threshold, similar to what is observed psychophysically as a function of polar angle, the computational observer model would require an additional 7 diopters of defocus. This is far higher than any plausible difference in defocus as a function of polar angle at 4.5°. Typically, defocus at 4.5° along the horizontal or vertical meridian is within ~0.2 diopters of defocus at the fovea (43, 44). The difference between the vertical and horizontal locations at 4.5° would be even less. Assuming a difference in defocus of 0.2 diopters, the optical quality would explain only about 3% of the effect of visual performance as a function of polar angle for this task.

### 2.4 The effect of cone density on orientation discrimination

The cone mosaic varies substantially with retinal location. As eccentricity increases, cone diameter increases, as does spacing between the cones, resulting in lower density. We used our computational observer model to quantify the extent to which variations in the cone mosaic could explain the changes in performance with polar angle. We simulated a large range of cone densities, spanning a range from about 3 times lower to 15 times greater than the typical density at 4.5° eccentricity (i.e., the retinal location of the simulated psychophysical experiment). As we varied the cone density, we also varied the cone size and spacing between cones according to the reported relation between density and coverage (40). The denser mosaics sample the stimulus more finely, with fewer absorptions per cone, because the cone area decreases it captures less photons. (**Fig 8A**).

**Fig 8.**
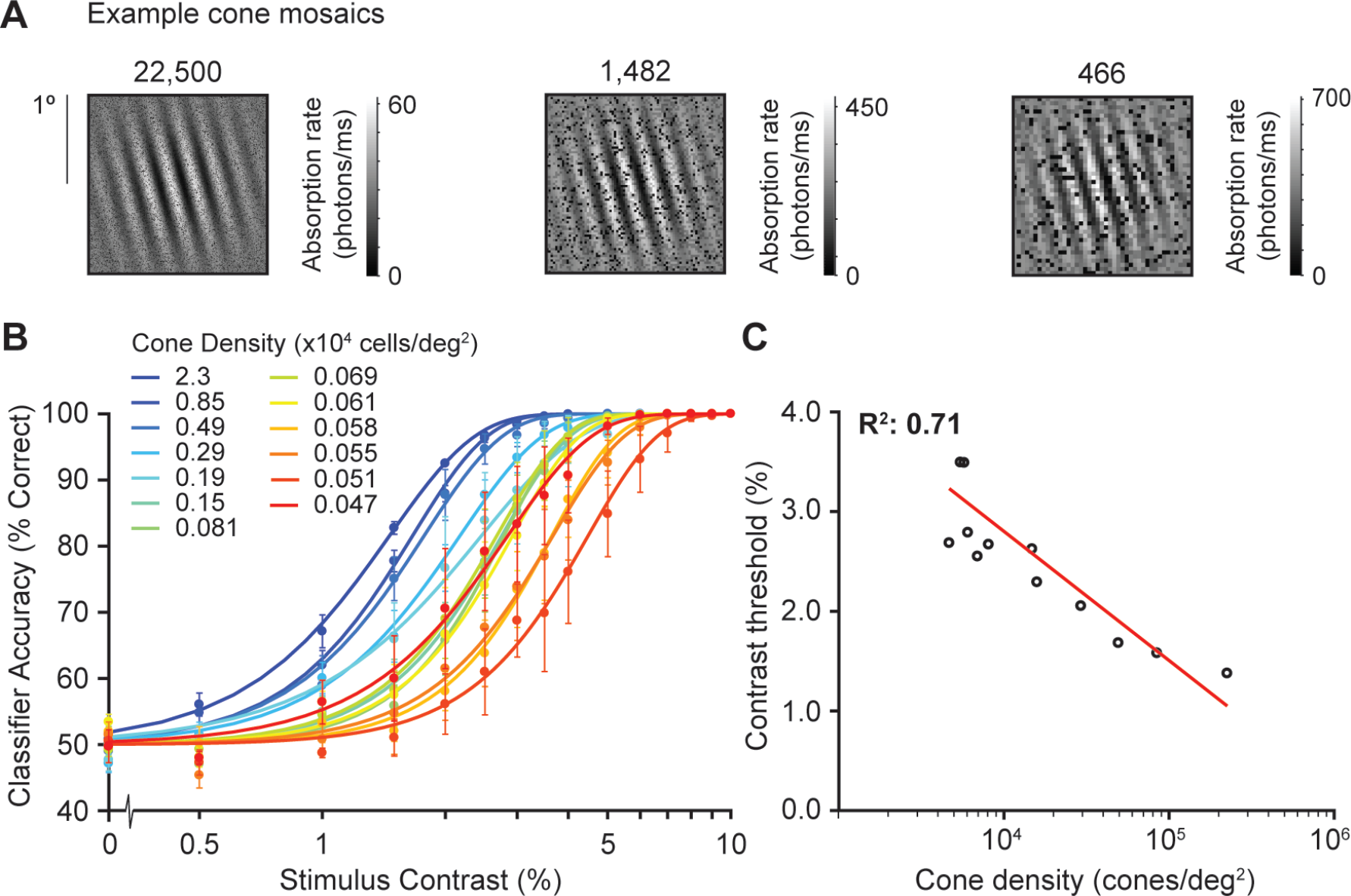
The effect of cone density on model performance. **(A)** Example cone mosaics simulated by our computational observer model. The highest tested cone density (22,500 cones/deg^2^) is equivalent to the cone density in the fovea (0°) whereas the lowest (466 cones/deg^2^) is equivalent to a cone density at 40° eccentricity, according to data from Curcio et al. (40). In our computational observer model, lower cone density implicitly results in larger absorption rates, because the cone area increases and therefore captures more photons. **(B)** Classifier accuracy for absorption rates as a function of stimulus contrast for 13 different cone density levels, each psychometric function is the average across performance of 2,000 trials per stimulus contrast (30,000 trials in total). Error bars represent standard error of the mean across 5 simulated experiments with 400 trials for each stimulus contrast. **(C)** Contrast thresholds from panel B as a function of log cone density (cones/deg^2^), fitted with a log-linear function.

Our computational observer model shows a decrease in contrast threshold as a function of cone density (**Fig 8B** **and C**). However, the effect is relatively small. For every 5-fold increase in cone density, the computational model contrast threshold reduces by 1 percentage point (e.g., from 4% to 3%). The meridional effect on human performance is ~4.4% (upper vertical meridian) vs 3.4% (horizontal). For cone density to account for this observed difference in human contrast thresholds, there would need to be about a 500% meridional difference in cone density. This is far greater than the 20-30% reported difference in cone density at 4.5° eccentricity (40, 41, 56). This indicates that, according to our computational observer model, cone density accounts for less than 10% of the differences in visual performance with polar angle on the orientation discrimination task reported by Cameron et al. (5).

## 3. Discussion

### 3.1 An explicit model is needed to link biological measurements with psychophysical performance

Our goal was to assess the degree to which front-end properties of the visual system explain well-established psychophysical performance differences around the visual field. In particular, we quantified the contribution of two factors in the eye – cone density and optical quality (defocus) – to contrast thresholds measured at different polar angle in an orientation discrimination task as reported by (5). These front-end factors have been reported to vary with polar angle, and in principle, the observed performance differences could be a consequence of the way the first stages of vision process images. For instance, cone density is higher on the horizontal meridian compared to the vertical meridian (up to 20° eccentricity (40, 41, 56)). Nonetheless, without a model to link these factors to performance, how much explanatory power they have cannot be assessed. We therefore developed a computational observer model to test these potential links. The underling software we used, ISETBIO, has recently been used to model a number of basic psychophysical tasks, including contrast sensitivity (46), Vernier acuity (57), illumination discrimination (58), color perception (59), chromatic aberration (60), visual perception with retinal prosthesis (61), and spatial summation in Ricco’s area (62).

### 3.2 Optics and cone density can explain only a small part of performance fields

Although cone density along the cardinal meridians correlates with behavior, our model showed that this correlation has little explanatory power: Differences in cone density can only account for a small fraction of the variation in visual performance as a function of meridian. Similarly, variation in optical quality within a plausible biological range has only a very small effect on contrast thresholds in the model of our task. Our observer model puts a ceiling on these two factors at less than 10% of the observed psychophysical effects. To fully explain these visual performance differences with polar angle, our computational model would require a difference of more than 7 diopters in defocus and a difference of more than 500% in cone density for the horizontal compared to the upper vertical meridian. Such large differences are far outside the range of plausible biological variation; defocus at 4.5° eccentricity is typically within 0.1-0.2 diopters of the fovea (44) and cone density at the horizontal meridian is ~20-30% more than the vertical at this eccentricity (40).

### 3.3 Downstream processing contributes to performance fields

The fact that neither optics nor the cone sampling array can explain more than a small fraction of the effect of polar angle on contrast thresholds indicates that downstream mechanisms must explain the majority of this effect.

A potential contributing factor is the spatial pooling of retinal ganglion cells. Like photoreceptors, midget retinal ganglion cells sample the visual field asymmetrically. For example, at 4.5° eccentricity, the density of midget retinal ganglion cells on the horizontal meridian is reported to be 1.4 times greater than on the vertical meridian (~1,330 vs ~950 cells/deg^2^ on the horizontal vs. inferior retina) (56, 63). This 40% meridional effect is larger than the 20-30% effect at the level of the cones, indicating that polar angle asymmetries in cone density are accentuated in further retinal processing. We have not included retinal ganglion cells in our model, but given our observer model with the cone array, we speculate that this further meridional difference in ganglion cell density will not be sufficient to explain the reported meridional psychophysical effects.

A second potential factor is visual cortex. Some aspects of performance fields manifest as amplitude differences in the BOLD fMRI signal in V1. Liu, Heeger and Carrasco (64) reported a 40% larger BOLD amplitude in V1 for stimuli on the lower than the upper vertical meridian. This asymmetry was found for high but now low spatial frequency stimuli, matching psychophysical results. They did not report differences between stimuli on the vertical versus horizontal meridians. Performance fields may also be reflected in the geometry of visual cortex. For example, a template of the V1 map fit to a population of 25 observers showed more cortical area devoted to the horizontal than the vertical meridian, although the authors acknowledged that this could be a fundamental fact about V1 or an artifact of the flattening process used in their analyses (65). This areal difference has been confirmed in an independent data set (66). These data also showed that population receptive fields (pRFs) in V1 and V2 are ~10% smaller when comparing horizontal to vertical quadrants. The geometry and the pRF size effects are complementary: greater area and smaller pRFs along the horizontal meridian are both consistent with this part of visual cortex analyzing the visual field in greater detail. However, there are also psychophysical differences between the upper and the lower vertical meridian (2, 6, 7), for which no pRF differences were reported. Moreover, there is not an explicit model to link differences in pRF properties directly to performance on a particular psychophysical task. Hence it is unknown whether and how these cortical differences would account for the observed behavioral patterns.

In addition to factors in early visual cortex, cognitive factors will also be important to consider in developing a full understanding of visual performance across polar angles. Exogenous covert visual attention does not compensate for discriminability differences across polar angles (4–6, 19), but endogenous covert attention may do so. We are currently investigating this possibility.

### 3.4 Limitations of the model

Our goal in building a computational observer model was to explicitly link known facts about the biology of the visual system with psychophysical performance. The value of the model is evidenced by the difference in the inference one might have drawn from a purely correlational approach (performance is best where cone density is highest) and the inference drawn from the model (little relation between cone density and performance). Nonetheless, all models are simplifications, and ours is no exception.

First, our model contained only one eye, whereas most of the psychophysical evidence in support of performance fields comes from binocular experiments. There have, however, also been some monocular experiments that vary stimulus polar angle, and these experiments confirm differences in performance across polar angle and show a similar magnitude of the effect (4, 8). Hence this limitation is unlikely to affect our conclusions.

Second, we modeled the cone mosaic as a rectangular patch with uniform density for each simulation, whereas the photoreceptors in human retina are organized in a hexagonal grid with a gradual change in density as a function of eccentricity. The uniformly spaced rectangular grid was implemented to save computational resources. The difference between an eccentricity-dependent mosaic and a uniform mosaic can be important for modeling performance near the fovea (59), as density declines rapidly over a short distance (40). However, further in the periphery, the density changes are modest across a small patch. And given that our model showed that very large differences in the cone array were needed to explain variation in psychophysical performance, it is unlikely that using a hexagonal, eccentricity-dependent array would have altered our conclusions.

Third, we did not model differences in photopigment density or macular pigment density as a function of retinal position. Pigment density has an effect on wavelength sensitivity and overall efficiency (48). Although our model did not vary pigment density, it did include position-dependent efficiency, implemented by varying the cone coverage, which ranged from close to 1 (no gaps between cones) near the fovea to ~0.25 in the far periphery. Hence, additional variation in efficiency arising from pigment density would be unlikely to have a substantial impact on model performance. Moreover macular pigment density does not vary systematically with polar angle at iso-eccentric locations (67).

Finally, our computational model only deals with visual processes up to photon absorptions by the cones. Processes up to this point, optics, photon noise, and cone sampling, are well characterized and can be accurately modeled. In future work, we will build on our computational observer model to investigate the contribution of downstream factors, such as post-receptor retinal circuitry and pooling of signals by retinal ganglion cells and visual cortex.

### 3.5 The inference engine

The performance of a classifier depends, in part, on how much knowledge of the task the classifier has access to. Our observer model had far less information than an ideal observer model. By definition, the ideal observer model has complete knowledge about the stimuli and sets an upper limit on performance (68, 69). When ideal observer models are applied to very early signals in the visual system such as cone responses, they typically outperform human observers by a large margin, e.g., by a factor of 10 or more (49, 70). The incomplete knowledge in our observer model led to poorer performance than would be obtained by an ideal observer model, and similar performance to human observers (~2-4% contrast thresholds in our task). Moreover, although some observers may have some (explicit or implicit) knowledge of performance differences across eccentricity, they do not have knowledge of differences across iso-eccentric locations. In the task reported by (5), two of the observers were the authors (and trained psychophysical observers) and one observer was naïve. This suggests that knowledge of the phenomenon does not alter the pattern of performance.

Our inference engine has two types of knowledge about the task, one more general about visual processing and one more specific to the particular experiment we simulated. The general (and implicit) knowledge arises from transforming the 2D time-varying cone absorption images to amplitude spectra. Transforming the data in this way effectively gives the observer model knowledge that spatial frequency and orientation (but not phase) might be relevant for the task. This transform does not indicate which spatial frequencies or orientations are relevant. Although the visual system does not literally compute a Fourier transform of the cone responses, cells in visual cortex are tuned to orientation and spatial frequency in local patches of the image (71, 72), and complex cells in V1 are relatively insensitive to phase (73, 74). Hence the implicit knowledge we provide to the classifier via transform to the amplitude spectra is an approximation to general processing strategies in the visual system, rather than specific knowledge about our particular task. Pilot simulations in which the classifier operated directly on the absorption images resulted in near-chance performance. This is expected, because the phase randomization of the stimuli causes the number of absorptions for any particular cone to be uninformative as to the stimulus orientation.

More specific knowledge in the computational observer model comes from the training trials, which are used to learn the best linear separation (hyperplane) between the two classes. The plane is defined by a weighted sum of the classifier inputs (amplitude spectra in our case), which can be thought of as an approximation to receptive field analysis by downstream neurons. The high weights learned by the classifier for this task correspond to oriented, band-pass filters, which match properties of the stimuli (**Fig 5**, panel 4). Because the model has incomplete knowledge, values far from the stimulus (very high or very low spatial frequency, and orientations far from the stimulus orientations) have non-zero weights, which are learned during training on a finite number of noisy trials.

## Conclusion

Overall, our model includes a relatively detailed, biologically plausible front-end, which incorporates realistic details about the optics, photon noise, small fixational eye movements, and wavelength- and position-sampling by photoreceptors. This front-end processing was combined with a linear classifier that performs at levels comparable to the human without providing explicit knowledge about the tasks. Future work will incorporate more biologically explicit models of downstream processing, including retinal and cortical circuitry. Such models are likely to reveal that later processing in the nervous system amplifies asymmetries in processing around the visual field that begin in the earliest stages of vision, and thus, to explain a larger portion of the psychophysical asymmetries found in many visual tasks.

## 4. Methods

### 4.1 Computational observer model software overview

The computational observer model relies on the publicly available, MATLAB-based Image Systems Engineering Toolbox for Biology (ISETBIO (45–47)), available at http://isetbio.org/. The ISETBIO toolbox incorporates the image formation process, wavelength-dependent filtering, optical quality, and the spatial arrangement and biophysical properties of cones. We used the ISETBIO toolbox for the core model architecture and supplemented it with experiment-specific custom MATLAB code. The experiment-specific code implements stimulus parameters matched to a prior psychophysical study (5), manipulation of biological parameters to assess their impact on performance, and a 2-AFC linear support vector machine classifier. In the interest of reproducible computational methods, the experiment-specific code, for both simulation and analysis, is publicly available via GitHub (http://github.com/isetbio/JWLOrientedGabor). In addition, the data structures created by the simulation and analyses are permanently archived on the Open Science Framework URL: https://osf.io/mygvu/.

### 4.2 Psychophysical experiment

Our simulations were created to match a previous psychophysical study (5). In that study, stimuli were achromatic oriented Gabor patches. The Gabors were comprised of harmonics of 4 cycles/°, windowed by a Gaussian with a standard deviation of 0.5°, presented at 4.5° eccentricity, at one of 8 locations equally spaced around the visual field (see also **Fig 1A**). Gabor patches were tilted either 15° clockwise or counter-clockwise from vertical, and presented for 54 ms on each trial. The contrast of the Gabor patches varied from trial to trial. The contrast levels were selected for each observer based on pre-experiment testing, and usually ranged from about 1% to 10% Michelson contrast using a method of constant stimuli. The observer’s task was to indicate the orientation of the Gabor stimulus relative to vertical (clockwise or counter-clockwise) with a button press. Data were analyzed by fitting a Weibull function to the mean performance (% correct) at each contrast level, independently at different locations around the visual field.

### 4.3 Stimuli (scene spectral radiance)

The observer model starts with a description of the stimulus, called a ‘scene’ in ISETBIO. The scene is defined by the spectral radiance at each location in space and time (the ‘light field’). The spectral radiance contained wavelengths ranging from 400-700 nm, discretized to 10 nm steps, with equal photons at each wavelength (3.8×10^15^ quanta/s/sr/nm/m^2^). The stimulus was discretized into 2-ms time steps and 1.8-arcminute spatial steps (32 samples per degree). The scene comprised Gabor stimuli with parameters described above (section 4.1.1), oriented either clockwise or counter-clockwise, represented within a field of view of 2° diameter, and presented for 54 ms per trial. The dimensions of the scene were therefore 64 × 64 × 31 × 28 (height × width × wavelength × time). Gabor patches varied in Michelson contrast between 0.05% and 10%. We also incorporated a stimulus with 0% contrast stimulus as a sanity check whether our model would perform at chance level. For all stimuli, the mean luminance was 100 cd/m^2^. Because photon noise and eye movement noise are added later (see sections 4.14 and 4.15), and because we do not model the scene before or after the stimulus onset/offset, the scene is in fact identical at all 28 time points.

Machine learning algorithms can exploit sources of information that a human observer would be unlikely to use. For example, if the value of a single image pixel happened to correlate with the stimulus class, a classifier could succeed based on only the value of this pixel. We wanted to prevent our classifier from succeeding in this way. In our simulations (unlike the Cameron et al. paper (5)), the phase of the Gabor patches was selected from two values 180° apart (φ = 90° and φ = 270°), randomized across trials. A 180° phase difference means that the two possible stimuli within a class were identical except for a sign reversal. As a result, the expected value of each pixel in each stimulus class was 0 (relative to the background). Similarly, the expected value of the cone absorption rates at each location on the retina within a stimulus class was 0 (relative to the background). Therefore, the linear classifier could not succeed using the absorption level from any single cone. We believe human observers do not perform the task this way either, hence randomizing the phase is likely to make the observer performance more similar to the human performance.

### 4.4 Optics (retinal irradiance)

The optics transform the scene into a retinal image. We first describe the optics used for the simulations in sections 2.2 and 2.4 (**Fig 6** and **Fig 8**). For these simulations, the optics were matched to a typical human eye with a 3-mm pupil (diameter) in focus at 550 nm using a statistical model of wavefront aberrations (54). This statistical model is based on measurements from healthy eyes of 100 observers (55), and described by a basis set of Zernike polynomials (75). The statistical model by Thibos contained the first 15 Zernike coefficients (Z0-Z14, using OSA standard indexing). The simulated human wavefront was used to construct a point spread function (PSF). This PSF was convolved with the scene at every time point to generate the retinal image. After this spatial blurring, the optical image was further transformed by spectral filtering (light absorption by inert pigments in the lens and macula), which primarily reduce the intensity of short-wavelength light. Finally, the optical images were padded by 0.25° on each side with the mean intensity at each wavelength. The padding is needed to handle eye movements, so that cones near the edge of the simulated retinal patch have a defined input even when these cones are moved outside the scene boundaries. The dimensions of the optical image are the same as the dimensions of the scene, except for the spatial padding: 80 × 80 × 31 × 28 (height × width × wavelength × time), which was discretized the same way as the scene.

To investigate the effect of optical quality on visual performance of our task, we systematically added further defocus to the model of human optics (section 2.3, **Fig 7**). We did this by increasing the Z4 Zernike coefficient (defocus) from 0-2 μm in steps of 0.25 μm (corresponding to 0-6.16 diopters for a 3-mm pupil), while keeping all other Zernike coefficients from Thibos’ statistical model unchanged. Note that using a defocus coefficient of 0 does not result in perfect diffraction limited optics, given that the other aberrations are still non-zero. We manipulated defocus rather than all the higher-order aberrations because at the stimulus eccentricity we simulated (4.5°), defocus is the largest contributor to optical quality (44).

### 4.5 Cone mosaic: spatial sampling and isomerization

We constructed the cone mosaic as a uniformly spaced rectangular patch with a field of view matched to the stimulus (2×2°). Each cone mosaic contained a random distribution of L-, M- and S-cones with a ratio of 0.6:0.3:0.1. We used the Stockman-Sharp (76) functions to estimate cone photopigment spectral sensitivity, assuming 50% optical density for L- and M-cones, and 40% for S-cones. Peak efficiency was assumed to be equal for each cone class, 66.67% multiplied by the retinal coverage (the fraction of local retina occupied by cones).

For the simulations in sections 2.2 and 2.3 (**Fig 6** and **Fig 8**), the cone density was 1,560 cells/deg^2^, approximately matched to the density at 4.5° on the horizontal retina as reported by Curcio et al. (40). This results in an array of 79 × 79 cones for our 2° patch. The positions of the L-, M-, and S-cones were randomized within the array (but held to fixed ratio). For these simulations, we assumed a coverage proportion of 0.49, meaning that the cone inner segments sampled from about half of the optical image, and missed about half due to the spaces between cones. A coverage of less than 1 acts like a reduction in efficiency, since photons are lost to the gaps between cones. In general, cone coverage decreases with eccentricity as the density of rods increases, filling the spaces between cones.

In one set of experiments, we systematically varied cone density (Results section 2.4, **Fig 8**), spanning 22,500 to 466 cones/deg^2^ (corresponding to cone arrays ranging from 297 × 297 to 43 × 43). For each cone density, we determined an equivalent eccentricity based on the relation between eccentricity and density on the nasal meridian from Curcio et al. (40). We then adjusted the cone coverage according to this eccentricity, assuming that coverage declines exponentially as a function of eccentricity, from 1 (fovea, no gaps between cones) to 0.25 at 40°. This approximation is similar to that used by Banks et al. (49), which was based on data from Curcio et al. (40).

The number of absorptions was computed for each cone in two steps. First, the noiseless number of absorptions was computed by multiplying the appropriate cone sensitivity function (L, M, or S) by the corresponding location in the optical image (hyperspectral), and scaling this value by the peak efficiency (66.67%). The cone coverage was accounted for by only sampling the optical image at the locations within the cone inner segments. Second, the noiseless values were converted to noisy samples by assuming a Poisson distribution.

The dimensions of the cone array absorptions were 79 × 79 × 28 (rows × columns × time) for the simulations in sections 2.2 and 2.3 (**Fig 6** and **Fig 8**). When the cone density varied (section 2.4, **Fig 7**), the first two dimensions of the cone array size also changed.

### 4.6 Eye movements

We added small fixational eye movements (drift and microsaccades), before computing the isomerization rate for each cone at each time sample. The ISETBIO toolbox provides code that generates eye movement samples based on a Mergenthaler and Engbert’s drift model (50) and microsaccade statistics reported by Martinez-Conde et al. (51, 52).

The drift model computes eye movement paths for a single trial with modified Brownian motion process. The eye movement paths were generated in units of arc minutes and then converted to discrete cone shifts in the horizontal and vertical direction. If the amplitude of an eye movement was smaller than the distance between two cones, the displacement was accumulated over multiple time samples, until the threshold was reached, before a new shift was added to the eye movement path.

The drift model was implemented by adding a displacement vector to the current position at each time point. The displacement vector was determined by combining 3 inputs: 2D Gaussian noise, an autoregressive term for persistent dynamics at short time scales, and a delayed negative feedback for antipersistent dynamics at longer time scales. The parameters we used for this model were the ISETIO defaults, which contained a horizontal and vertical delay defined as X = 0.07 s and Y = 0.04 s, feedback steepness of 1.1, and feedback gain of 0.15. The control function had a mean of 0 and standard deviation of 0.075 and the gamma parameter was set to 0.25. The mean noise position and standard deviation were set to 0 and 0.35, respectively. Before computing the velocity of a drift period, the drift model applied a temporal smoothing filter to the eye movement paths using a 3^rd^ order Savitzky-Golay filter over a velocity interval of 41 ms.

For periods where the drift was stabilized, the eye movement code checked for microsaccade jumps to the eye movement path. Whether or not a microsaccade was added depended on when the last microsaccade was. In our experiment, we used the ISETBIO default where the interval between microsaccades followed a gamma function with a mean of 450 ms, with a minimum duration of 2 ms. A microsaccade was defined as a vector with a mean amplitude of a microsaccade was 8 arc minutes. Each vector contained an additional endpoint jitter of 0.3 arc min in length and 15° in direction. The microsaccade jumps were either ‘corrective’ (towards the center of the mosaic) or ‘random’ (any direction). The microsaccade mean speed was defined as 39°/s, with a standard deviation of 2°/s. Given that the defined interval between microsaccades was long compared to the stimulus duration (54 ms), most trials did not contain microsaccades.

A 216 ms warmup period was implemented before the trials began. Eye movements during this period affected the eye position at the start of the trial but were not otherwise included in the analysis.

### 4.7 Simulated Experiments and Behavioral Inference

A simulated experiment comprised 6,000 trials, with 400 trials at each of 15 contrasts. The 400 trials per contrast level included 200 clockwise and 200 counterclockwise stimuli, each of which was further subdivided into 100 trials at each of 2 phases. The data from a single contrast level within a single experiment were represented as a 4D array (*m* rows × *m* columns × 28 time-points × 400 trials), in which *m* is the number of cones along one side of the retinal patch (79 in the experiments for **Fig 6** and **Fig 7**, variable for the experiments in **Fig 8**).

Within the 6,000 trials of an experiment, all parameters other than the stimulus orientation (clockwise or counterclockwise) and phase (90° or 270°) were held constant, including the spatial distribution of L-, M-, and S-cones, the optics, the cone density, the cone coverage, and the presence or absence of fixational eye movements. Each simulated experiment was repeated 5 times, so that a single psychometric function summarized 30,000 trials. The arrangement of L-, M-, and S-cones was regenerated randomly for each of the 5 repeated experiments. Error bars in **Fig 6-8** indicate standard errors of the mean across the 5 experiments.

Classification (clockwise vs. counter-clockwise) via cross-validation was performed separately for each stimulus contrast level (set of 400 trials) in each experiment as follows. First, each *m* × *m* image of cone absorptions was transformed into an *m* × *m* amplitude spectrum using the 2D fast Fourier transform and discarding the phase information. This left the dimensionality unchanged (*m* rows × *m* columns × 28 time-points × 400 trials within a single contrast level). Second, the amplitudes were concatenated across space and time into a 2D matrix (400 trials × 28*m*2 values per trial). A 400-element vector labeled the trials by stimulus class (1 for clockwise and −1 for counter-clockwise). Third, the 2D matrix and 400-element vector with labels was used for training and testing a linear support vector machine (SVM) classifier on the amplitude images using MATLAB’s *fitcsvm* with 10-fold cross-validation, kernel function set to ‘linear’, and the built-in standardization option (to z-score each row of the data matrix). The learned classifier weights represented the best linear separation (hyperplane) between the two stimulus classes. With these trained weights, the classifier predicted the stimulus class label for the left out trials in a given data fold. We used MATLAB’s *kfoldLoss* function to average the accuracy across the 10-folds, which yielded one accuracy measure (% correct) per contrast level per experiment.

### 4.8 Quantifying the contribution of cone density and optics on the computational observer performance

To quantify the contribution of a given factor in the eye, we averaged the classifier accuracy for each stimulus contrast level across the 5 experiments to fit with a Weibull function (Equation 1). This resulted in full psychometric functions for each cone density and optical quality level. We calculated error bars for each contrast level as the standard error of the mean across the 5 iterations. For each psychometric function, we defined the contrast threshold as the power of 1 over the slope of the Weibull function β, in our case β = 3, of the performance level expected at chance (0.5 for a 2-AFC task, see Equation 2). This results in a threshold of ~80% (0.5^1/3^ = 0.7937).

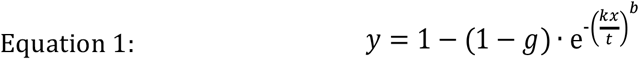

Where *g* is the performance expected at chance (0.5), *t* is the threshold, *b* is the slope of the Weibull function, and k is defined as:

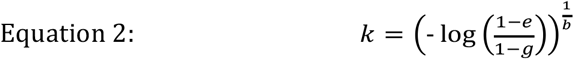

The contrast thresholds, *t*, were summarized as a linear function of defocus (**Fig 7**) or an exponential function of cone density (**Fig 8**, represented as a straight-line on a semi-log axis). These linear or log-linear fits enabled us to compute the change in cone density or the change in defocus needed to achieve a 1% increase in contrast threshold – similar to the meridional effect observed in human performance (~4.4% at the upper vertical meridian vs ~3.4% at the horizontal meridian as seen in (5)).

## Acknowledgments

This research was supported by the US National Eye Institute R01-EY027401 (M.C. and J.W.). We would like to thank Brian Wandell, David Brainard and Nicholas Cottaris for their advice and encouragement. We thank Pablo Artal for kindly providing data on human optical quality.

## References

1. Rovamo J, Virsu V, Nasanen R. Cortical magnification factor predicts the photopic contrast sensitivity of peripheral vision. Nature. 1978;271(5640):54–6.

2. Abrams J, Nizam A, Carrasco M. Isoeccentric locations are not equivalent: the extent of the vertical meridian asymmetry. Vision Res. 2012;52(1):70–8.

3. Strasburger H, Rentschler I, Juttner M. Peripheral vision and pattern recognition: a review. J Vis. 2011;11(5):13.

4. Carrasco M, Talgar CP, Cameron EL. Characterising visual performance fields: effects of transient covert attention, spatial frequency, eccentricity, task and set size. Spat Vis. 2001;15(1):61–75.

5. Cameron EL, Tai JC, Carrasco M. Covert attention affects the psychometric function of contrast sensitivity. Vision Res. 2002;42:949–67.

6. Talgar CP, Carrasco M. Vertical meridian asymmetry in spatial resolution: visual and attentional factors. Psychon Bull Rev. 2002;9(4):714–22.

7. Fuller S, Rodriguez RZ, Carrasco M. Apparent contrast differs across the vertical meridian: visual and attentional factors. J Vis. 2008;8(1):16 1-.

8. Corbett JE, Carrasco M. Visual performance fields: frames of reference. PLoS One. 2011;6(9):e24470.

9. Rovamo J, Virsu V. An estimation and application of the human cortical magnification factor. Experimental brain research. 1979;37:495–510.

10. Virsu V, Rovamo J. Visual resolution, contrast sensitivity, and the cortical magnification factor. Experimental brain research. 1979;37(3):475–94.

11. Rijsdijk JP, Kroon JN, van der Wildt GJ. Contrast sensitivity as a function of position on retina. Vision Res. 1980;20:235–41.

12. Robson JG, Graham N. Probability summation and regional variation in contrast sensitivity across the visual field. Vision Res. 1981;21(3):409–18.

13. Lundh BL, Lennerstrand G, Derefeldt G. Central and peripheral normal contrast sensitivity for static and dynamic sinusoidal gratings. Acta Ophthalmol (Copenh). 1983;61(2):171–82.

14. Regan D, Beverley KI. Visual fields for frontal plane motion and for changing size. Vision Res. 1983;23(7):673–6.

15. Skrandies W. Human contrast sensitivity: regional retinal differences. Hum Neurobiol. 1985;4(2):97–9.

16. Anderson RS, Wilkinson MO, Thibos LN. Psychophysical localization of the human visual streak. Optom Vis Sci. 1992;69(3):171–4.

17. Mackeben M. Sustained focal attention and peripheral letter recognition. Spat Vis. 1999;12(1):51–72.

18. Altpeter E, Mackeben M, Trauzettel-Klosinski S. The importance of sustained attention for patients with maculopathies. Vision Res. 2000;40(10-12):1539–47.

19. Carrasco M, Williams PE, Yeshurun Y. Covert attention increases spatial resolution with or without masks: support for signal enhancement. J Vis. 2002;2(6):467–79.

20. Seiple W, Holopigian K, Szlyk JP, Wu C. Multidimensional visual field maps: relationships among local psychophysical and local electrophysiological measures. J Rehabil Res Dev. 2004;41(3A):359–72.

21. Silva MF, Maia-Lopes S, Mateus C, Guerreiro M, Sampaio J, Faria P, et al. Retinal and cortical patterns of spatial anisotropy in contrast sensitivity tasks. Vision Res. 2008;48(1):127–35.

22. Silva MF, Mateus C, Reis A, Nunes S, Fonseca P, Castelo-Branco M. Asymmetry of visual sensory mechanisms: electrophysiological, structural, and psychophysical evidences. J Vis. 2010;10(6):26.

23. Silva MF, d’Almeida OC, Oliveiros B, Mateus C, Castelo-Branco M. Development and aging of visual hemifield asymmetries in contrast sensitivity. J Vis. 2014;14(12).

24. Carrasco M, Giordano AM, McElree B. Temporal performance fields: visual and attentional factors. Vision Res. 2004;44(12):1351–65.

25. Chaikin JD, Corbin HH, Volkmann J. Mapping a field of short-time visual search. Science. 1962;138(3547):1327–8.

26. Kristjansson A, Sigurdardottir HM. On the benefits of transient attention across the visual field. Perception. 2008;37(5):747–64.

27. Krose BJ, Julesz B. The control and speed of shifts of attention. Vision Res. 1989;29(11):1607–19.

28. Najemnik J, Geisler WS. Eye movement statistics in humans are consistent with an optimal search strategy. J Vis. 2008;8(3):4 1–14.

29. Najemnik J, Geisler WS. Simple summation rule for optimal fixation selection in visual search. Vision Res. 2009;49(10):1286–94.

30. Pretorius LL, Hanekom JJ. An accurate method for determining the conspicuity area associated with visual targets. Hum Factors. 2006;48(4):774–84.

31. Rezec AA, Dobkins KR. Attentional weighting: a possible account of visual field asymmetries in visual search? Spat Vis. 2004;17(4-5):269—93.

32. Fortenbaugh FC, Silver MA, Robertson LC. Individual differences in visual field shape modulate the effects of attention on the lower visual field advantage in crowding. J Vis. 2015;15(2).

33. He S, Cavanagh P, Intriligator J. Attentional resolution and the locus of visual awareness. Nature. 1996;383(6598):334–7.

34. Toet A, Levi DM. The two-dimensional shape of spatial interaction zones in the parafovea. Vision Res. 1992;32(7):1349–57.

35. Fuller S, Carrasco M. Perceptual consequences of visual performance fields: the case of the line motion illusion. J Vis. 2009;9(4):13 1–7.

36. Montaser-Kouhsari L, Carrasco M. Perceptual asymmetries are preserved in short-term memory tasks. Attention, perception & psychophysics. 2009;71(8):1782–92.

37. Edgar GK, Smith AT. Hemifield differences in perceived spatial frequency. Perception. 1990;19(6):759–66.

38. Rodieck RW. The First Steps in Seeing. Oxford: Oxford University Press; 1998.

39. Wandell BA. Foundations of Vision: Sinauer Associates; 1995.

40. Curcio CA, Sloan KR, Kalina RE, Hendrickson AE. Human photoreceptor topography. The Journal of comparative neurology. 1990;292:497–523.

41. Song H, Chui TY, Zhong Z, Elsner AE, Burns SA. Variation of cone photoreceptor packing density with retinal eccentricity and age. Invest Ophthalmol Vis Sci. 2011;52(10):7376–84.

42. Artal P. Image Formation in the Living Human Eye. Annu Rev Vis Sci. 2015;1:1–17.

43. Polans J, Jaeken B, McNabb RP, Artal P, Izatt JA. Wide-field optical model of the human eye with asymmetrically tilted and decentered lens that reproduces measured ocular aberrations. Optica. 2015;2(2):124.

44. Jaeken B, Artal P. Optical quality of emmetropic and myopic eyes in the periphery measured with high-angular resolution. Invest Ophthalmol Vis Sci. 2012;53(7):3405–13.

45. Farrell JE, Winawer J, Brainard DH, Wandell B. 27.2: Distinguished Paper: Modeling visible differences: The computational observer model. SID Symposium Digest of Technical Papers 2014. p. 352–6.

46. Cottaris N, Jiang H, Ding X, Wandell B, Brainard DH. A computational observer model of spatial contrast sensitivity: Effects of wavefront-based optics, cone mosaic structure, and inference engine. bioRxiv. 2018.

47. Brainard DH, Jiang H, Cottaris NP, Rieke F, Chichilnisky EJ, Farrell JE, et al., editors. ISETBIO: Computational tools for modeling early human vision. Imaging and Applied Optics 2015; 2015 2015/06/07; Arlington, Virginia: Optical Society of America.

48. Wyszecki G, Stiles WS. Color science. New York: Wiley; 1982.

49. Banks MS, Sekuler AB, Anderson SJ. Peripheral spatial vision: limits imposed by optics, photoreceptors, and receptor pooling. J Opt Soc Am A. 1991;8(11):1775–87.

50. Mergenthaler K, Engbert R. Modeling the control of fixational eye movements with neurophysiological delays. Phys Rev Lett. 2007;98(13):138104.

51. Martinez-Conde S, Macknik SL, Hubel DH. The role of fixational eye movements in visual perception. Nature reviews Neuroscience. 2004;5(3):229–40.

52. Martinez-Conde S, Macknik SL, Troncoso XG, Hubel DH. Microsaccades: a neurophysiological analysis. Trends Neurosci. 2009;32(9):463–75.

53. Villegas EA, Alcon E, Artal P. Optical quality of the eye in subjects with normal and excellent visual acuity. Invest Ophthalmol Vis Sci. 2008;49(10):4688–96.

54. Thibos LN. Retinal image quality for virtual eyes generated by a statistical model of ocular wavefront aberrations. Ophthalmic Physiol Opt. 2009;29(3):288–91.

55. Thibos LN, Hong X, Bradley A, Cheng X. Statistical variation of aberration structure and image quality in a normal population of healthy eyes. J Opt Soc Am A. 2002;19(12):2329–48.

56. Curcio CA, Sloan KR, Packer O, Hendrickson AE, Kalina RE. Distribution of cones in human and monkey retina: Individual variability and radial asymmetry. Science. 1987;236(4801):579–82.

57. Jiang H, Cottaris N, Golden J, Brainard D, Farrell J, Wandell B, editors. Simulating retinal encoding: facors influencing vernier acuity. Proceedings of Electronic Imaging; 2017; Burlingame, CA.

58. Ding X, Radonjić A, Cottaris NP, Jiang H, Wandell BA, Brainard D. Computational-Observer analysis of illumination discrimination. 2018.

59. Brainard DH, Cottaris NP, Radonjic A. The perception of color and material in naturalistic tasks. BioRXiv. 2018.

60. Wang X, Pedersen M, Tomas JB. The influence of chromatic aberration on demosaicking. BioRXiv. 2014.

61. Golden JR, Erickson-Davis C, Cottaris NP, Parthasarathy N, Rieke F, Brainard DH, et al. Simulation of visual perception and learning with a retinal prosthesis. BioRXiv. 2018.

62. Tuten WS, Cooper RF, Tiruveedhula P, Dubra A, Roorda A, Cottaris NP, et al. Spatial summation in the human fovea: the effect of optical aberrations and fixational eye movements. BioRXiv. 2018.

63. Watson AB. A formula for human retinal ganglion cell receptive field density as a function of visual field location. J Vis. 2014;14(7).

64. Liu T, Heeger DJ, Carrasco M. Neural correlates of the visual vertical meridian asymmetry. J Vis. 2006;6(11):1294–306.

65. Benson NC, Butt OH, Datta R, Radoeva PD, Brainard DH, Aguirre GK. The retinotopic organization of striate cortex is well predicted by surface topology. Current biology: CB. 2012;22(21):2081–5.

66. Silva MF, Brascamp JW, Ferreira S, Castelo-Branco M, Dumoulin SO, Harvey BM. Radial asymmetries in population receptive field size and cortical magnification factor in early visual cortex. NeuroImage. 2018;167:41–52.

67. Robson AG, Moreland JD, Pauleikhoff D, Morrissey T, Holder GE, Fitzke FW, et al. Macular pigment density and distribution: comparison of fundus autofluorescence with minimum motion photometry. Vision research. 2003;43(16):1765–75.

68. Green DM, Swets JA. Signal detection theory and psychophysics. New York,: Wiley; 1966. xi, 455 p. p.

69. Geisler WS. Contributions of ideal observer theory to vision research. Vision research. 2011;51(7):771–81.

70. Banks MS, Geisler WS, Bennett PJ. The physical limits of grating visibility. Vision research. 1987;27(11):1915–24.

71. Blakemore C, Campbell FW. On the existence of neurones in the human visual system selectively sensitive to the orientation and size of retinal images. J Physiol. 1969;203(1):237–60.

72. Carandini M, Demb JB, Mante V, Tolhurst DJ, Dan Y, Olshausen BA, et al. Do we know what the early visual system does? J Neurosci. 2005;25(46):10577–97.

73. Hubel DH, Wiesel TN. Receptive fields, binocular interaction and functional architecture in the cat’s visual cortex. J Physiol. 1962;160:106–54.

74. Rust NC, Schwartz O, Movshon JA, Simoncelli EP. Spatiotemporal elements of macaque v1 receptive fields. Neuron. 2005;46(6):945–56.

75. Zernike F. Diffraction theory of the knife-edge test and its improved form, the phase-contrast method. Monthly Notices of the Royal Astronomical Society. 1934; 94(377-384).

76. Stockman A, Sharpe LT. The spectral sensitivities of the middle- and long-wavelength-sensitive cones derived from measurements in observers of known genotype. Vision Res. 2000;40(13):1711–37.

